# Release strategies affect the freshwater residence and survival of hatchery-reared juvenile Chinook Salmon

**DOI:** 10.1101/2024.12.20.629694

**Authors:** Thomas J. B. Balfour, David J. A. Hurwitz, Jamieson B. Atkinson, Eduardo G. Martins

## Abstract

**Objective:** This study investigated the effects of size, location and timing of release on the freshwater residence and survival hatchery-reared Chinook Salmon (*Oncorhynchus tshawytscha*) in the Toquaht River, BC.

**Methods:** Juvenile salmon were PIT tagged and released on three separate dates (May 23, June 9 and 19, 2021) and three distinct locations within the river (lower river [below lake], lake, and upper river [above lake]). Fish were detected near the river mouth using a PIT array, and detection data were analyzed with an integrated model of freshwater residence and capture-recapture using Bayesian inference.

**Result:** The median duration of freshwater residence was 13.8 days and was longer for fish released in the lake and upper river earlier in the study. The median survival probability was 0.35 and it was higher for fish released in the lake and upper river later in the study. Both freshwater residence time and survival declined with fish size at release.

**Conclusion:** These findings highlight that hatchery release strategies can significantly influence survival and freshwater residence times, underscoring the need for adaptable management practices.

**Impact statement:** Juvenile Chinook Salmon survival during freshwater residence and outmigration is strongly influenced by hatchery release strategies, thus optimizing these strategies is key for hatcheries to support the recovery of salmon populations.

## INTRODUCTION

It is broadly accepted that the rates at which Pacific salmon survive and return to spawn are determined by a complex web of interlinked factors over a wide range of spatial and temporal scales (Beamish et al. 2003; Ruff et al. 2017; Dorner et al. 2018). A core challenge in understanding how salmon respond to environmental changes is separating the influence of local from regional and global factors shared among populations (Beamish and Mahnken 2001; Rogers and Schindler 2011; Dorner et al. 2018). Historically, it has been difficult to determine rates of early marine mortality independently from mortality experienced during the juvenile freshwater rearing and outmigration phases (Michel 2019). However, the proliferation of small acoustic telemetry and passive integrated transponder (PIT) tags has allowed for investigations into the survival and behavior of salmon during discrete life stages (e.g. smolt outmigration) at a much finer scale than previously possible (Hammer and Blankenship 2001).

Recent studies using electronic tags on juvenile salmon have revealed outmigration survival probabilities ranging from very low (0.01) to moderate (0.50) and a large degree of variation across years (Chittenden et al. 2010; Aarestrup et al. 2014; Clark et al. 2016; Rechisky et al. 2019; Notch et al. 2020). Furthermore, the low survival estimates revealed by some studies suggest that outmigration success may have a larger impact on the abundance of Pacific salmon populations than previously thought, and in some cases may be the dominant period in determining adult returns (Chittenden et al. 2010; Michel 2019). Therefore, increasing our understanding of the freshwater outmigration period is an important step in making effective management decisions, and increasing the investment in outmigration studies across a wider array of salmon populations and river systems should be a top priority (James 2021).

Of the five species of Pacific salmon, Chinook Salmon *(Oncorhynchus tshawytscha*) are the largest, least numerous (Ohlberger et al. 2016), and one of the most important to the economies, cultures, and ecosystems within their range (Riddell et al. 2018). On the West Coast of Vancouver Island (WCVI) in British Columbia, Canada, Chinook Salmon are known to be predominantly ocean-type and occur in over 100 rivers (Fisheries and Oceans Canada 2014). However, total adult returns in this region have declined since the 1990s with little evidence of recovery (Brown et al. 2019). In 2020, the Committee on the Status of Endangered Wildlife in Canada (COSEWIC) recommended that WCVI Chinook Salmon (designatable units 24, 25 and 26) be listed as “threatened” under the Species at Risk Act (COSEWIC 2020). The COSEWIC assessment identified issues related to freshwater ecosystems, marine harvest, climate change, and hatchery releases as the leading factors in this decline. Recent reviews have highlighted significant data gaps related to understanding the effectiveness and risks of hatchery programs (James et al. 2023; Riddell et al. 2024). This emphasizes the need for further research on hatchery practices at the program level to inform hatchery reform, which has been highlighted as a core component of WCVI Chinook Salmon rebuilding (Irvine et al. 2024).

In North America, hatcheries have been used to augment Pacific Salmon populations since the late 1800’s and remain a core component of salmon management (Anderson et al. 2020; James et al. 2023). Although hatchery programs have generally succeeded in compensating for the high mortality experienced by wild salmon during the egg-to-fry or smolt stages, the efficacy of hatcheries as a rebuilding tool remains uncertain and controversial (Mobrand et al. 2005; Anderson et al. 2020; Riddell et al. 2024). Hatchery programs implement various strategies to optimize survival rates to achieve management goals and mitigate risks to wild populations (Beckman et al. 2017; Nelson et al. 2019; James et al. 2023). For example, hatchery managers can control rearing conditions such as: water source, rearing containers, density, and feeding schedules, as well as release strategies, which include fish size at release, as well as timing, and location of release. However, their effectiveness can vary greatly among hatcheries and years, necessitating studying rearing and release strategies on a program-by-program basis (Anderson et al. 2020; James et al. 2023).

In this study, we used PIT telemetry and integrated statistical models to investigate how strategies related to size at release as well as release location and timing affect freshwater residence and survival of a hatchery-origin Chinook Salmon population in the Toquaht River on the West Coast of Vancouver Island. We were interested in testing the following hypotheses: i) residence time would be short (0.5–5 days) and survival moderate to high (> 0.5) due to the short migration distance and limited number of predators in the Toquaht River (Pellet 2019); ii) fish released later in the study would have shorter freshwater residence (as their release timing moves further from the natural outmigration period) and lower survival (due to low water levels that could increase predation risk); iii) fish released closer to the estuary would have shorter freshwater residence and higher survival due to the shorter outmigration distance; and that iv) smaller fish would reside longer in freshwater (due to slower migration rates and/or to continue growing) and have lower survival (due to higher vulnerability to predation) than larger fish.

## METHODS

### Study site

The study was conducted in the Toquaht River, which is a fourth order stream, located in northeastern Barkley Sound on the West Coast of Vancouver Island (49°2’15.31”N, 125°21’27.84”W) (Figure 1A). The Toquaht River is the most valued salmon river within the territory of the Toquaht Nation and historically supported populations of Chum Salmon *O. keta*, Chinook Salmon, Coho Salmon *O. kisutch*, Sockeye Salmon *O. nerka*, Steelhead and Rainbow Trout *O. mykiss*, Coastal Cutthroat Trout *O. clarkii-clarkii*, and Dolly Varden *Salvelinus malma* (Duncan et al. 1990). The Toquaht River watershed has seen a sharp decline in all fish populations since the mid-1980s (Fisheries and Oceans Canada 2024). In response to this decline, a community hatchery, run by the Thornton Creek Salmon Enhancement Society (TCES), implemented a Chinook Salmon enhancement program designed to stabilize and rebuild the dwindling population. Despite the high level of hatchery input, the current adult returns remain well below historic abundance (Fisheries and Oceans Canada 2024) and unlikely to be self-sustaining without continued hatchery intervention.

**FIGURE 1.**
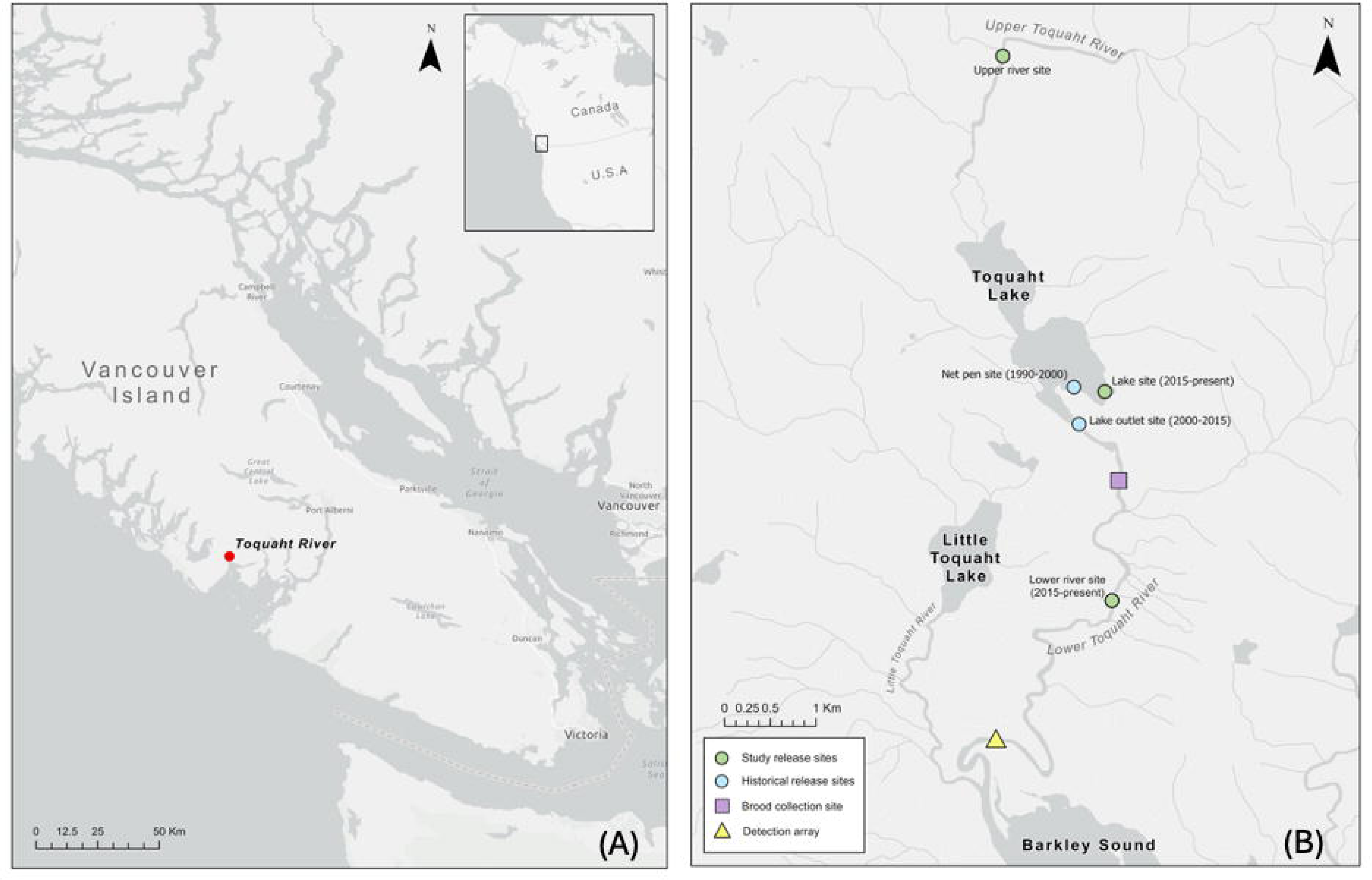
Maps showing the (A) location of the Toquaht River on Vancouver Island with an inset showing the study site location in Canada, and the (B) location of brood collection, release sites and the PIT array on the Toquaht River.

#### Brood collection and husbandry

On September 22^nd^ and October 6^th^ of 2020, Thornton Creek Hatchery staff collected Chinook Salmon brood stock by beach-seining and gillnetting from two locations in the mid section of the Toquaht River: Lucky Pool (49° 4’ 7.78”N, 125° 20’ 27.12”W) and Canyon Pool (49° 4’ 14.13”N, 125° 20’ 29.96”W) (Figure 1B). Brood stock fish were stripped of eggs or milt, then sampled streamside for tissue samples, scales, otoliths, and fork lengths. It is Thornton Creek Hatchery’s practice to use milt from more than one male to fertilize each female. This is done to mitigate for the possibility of an infertile male and to maximize genetic diversity in each batch (i.e. cohorts spawned on the same day that are comprised of fertilized eggs from one pair or several pairs of males and females; Table 1). All fish in a batch have the same accumulated thermal units (ATU) and are shocked at the eyed stage to separate viable from non-viable eggs. After hatching, fish were ponded in covered metal troughs with a freshwater flow-through system commonly used to rear hatchery salmon (each batch was reared in their own trough, but all were reared under the same conditions). Study fish were fed BioVita feed (Bio-Oregon, Longview, WA, USA) once daily, withholding food for one day every 7–10 days and increasing size of food to match growth rates of fish. Daily feed quantities were on based on BioVita guidelines, using rates between 0.016% to 0.025% of body weight per day throughout the rearing cycle based on water temperature and average body mass.

**TABLE 1.**
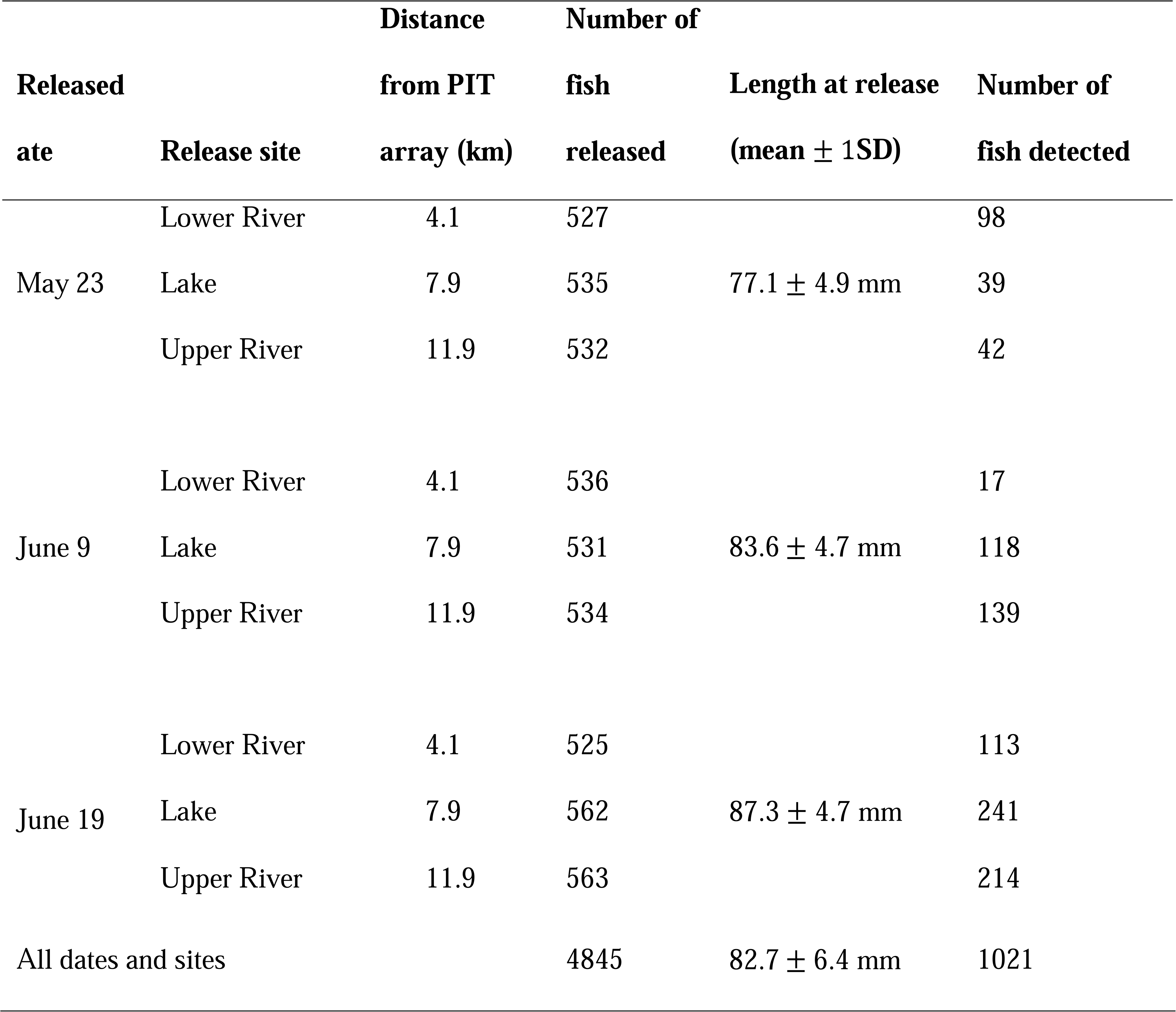
Summary of the number of fish released and detected as well as their length at release. Each release group is defined by a combination of release site and date. Three fish were excluded from the analyses because their length was not measured and were not used in the summaries shown in the table.

### Tagging and releases

A total of 5,000 fish were tagged once they surpassed a minimum tagging size threshold. Initially, we tagged approximately 1,000 fish using a 60 mm minimum fork length threshold, based on other regional PIT tag studies, but further switched to using a 69 mm minimum fork length threshold (Vollset et al. 2020). Fish were starved for 24 hours prior to tagging and then anesthetized using 0.1 g/L MS222. Anesthetized fish were implanted with a PIT tag (APT12 PIT 12.5 mm 134.2 kHz ISO FDX-B, Biomark, Boise, USA) into their abdominal cavity using MK25 PIT (Biomark, Boise, USA) tag implanters with preloaded needles. Tag implantation took place in a dedicated tagging trailer containing a custom designed flow-through table connected to the hatchery holding troughs so that tagged fish were immediately returned to their trough after receiving a tag. The number of fish selected for PIT tagging from each of the five batches was proportional to the number of fish in each batch (Table 1).

All tagged fish were monitored for up to 40 days post tagging. During this monitoring period, hatchery staff checked the rearing troughs twice daily for mortalities and collected rejected tags from the bottom of holding through using a 2.5 cm magnet screwed to the end of a 1 m long PVC pipe. We found 116 rejected tags and 165 mortalities of the 5,000 tagged fish, for a mortality rate of 3.1% and a tag shedding rate of 2.3%. Hatchery staff tagged 242 additional fish to compensate for tag loss and mortality. The two groups of fish tagged under different minimum size thresholds had negligible differences in mortality (2.9% and 3.4% for 60mm and 69mm threshold, respectively). During the pre-release scans (17-44 days after tagging), 4,848 fish still retained operational tags and were released. However, the length of three fish was not recorded and they were removed from the analyses, which were based on data of 4,845 fish.

Two to four days prior to release, each fish was anesthetized, PIT ID recorded, weighed, and measured for fork length and divided evenly in one of three holding troughs (Table 2). Tagged fish were released on May 23, June 9 and June 19, 2021 at all three release sites: the lower Toquaht River at the bridge (4.1 km upstream of the PIT array, see below), Toquaht Lake (7.9 km from the array), and the upper Toquaht River (11.9 km from the array) (Figure 1B). Compared to the standard release strategies utilized by the Thornton Creek Hatchery, the May 23 release date is earlier than usual, whereas the June 9 and June 19 dates fall within the typical release window. While the Lower River and Lake are frequently used as release sites, the Upper River had not been utilized in previous releases. Juveniles were transported to the release sites in two trucks equipped with 900 L aerated aluminium transfer tanks.

**TABLE 2.** Summary of observed freshwater residence times (days) by release site and date for fish that were detected at the PIT array. MAD: median absolute deviation.

| **Release site and date** | **Median** | **MAD** | **Min residence time (days)** | **Max residence time (days)** |
| --- | --- | --- | --- | --- |
| Lower River |  |  |  |  |
| May 23 | 0.7 | 0.1 | 0.5 | 34.6 |
| June 9 | 13.7 | 3.0 | 10.7 | 18.6 |
| June 19 | 2.6 | 2.9 | 0.7 | 17.6 |
| Lake |  |  |  |  |
| May 23 | 35.5 | 3.2 | 2.6 | 59.9 |
| June 9 | 18.5 | 2.9 | 10.6 | 42.6 |
| June 19 | 10.6 | 5.2 | 1.6 | 40.5 |
| Upper River |  |  |  |  |
| May 23 | 35.7 | 5.6 | 27.6 | 61.4 |
| June 9 | 18.7 | 3.2 | 10.7 | 48.6 |
| June 19 | 11.6 | 5.8 | 1.7 | 41.0 |

### PIT array placement and monitoring

The outmigration of PIT tagged Chinook Salmon was monitored by a PIT array made of two independent and identical detection antennas placed 100 m apart within the same shallow run in the lower Toquaht River. The array position was chosen to be as close to tidewater as possible while being broad and shallow to ensure best coverage of the migration pathway. Like most West Coast of Vancouver Island rivers, the Toquaht River receives a high volume of rainfall annually and in pulses that cause water levels to rise and drop quickly (Porter et al. 2017). The rapid rise and fall of the water levels meant each fish or group of fish would not always experience the same probability of detection when passing the PIT array. Indeed, our read-range tests revealed that the distance of a tag to the antenna, which would increase with rising water levels, would be an important factor influencing detection. Therefore, to account for variable detection probability caused by fluctuation between low and high water, water levels at the array location during the outmigration period were recorded using a HOBO U20L level logger (Onset, Bourne, USA). However, the water level logger failed and only collected data for the period between June 7 and 30, leaving missing records for most of the study period in 2021 (May 23 to August 29). Therefore, instead of using water level to account for variable detection probability in the model, we used rainfall data collected at a weather station located ∼16 km southwest of the study site (Kennedy Camp, 48°56’43.020” N 125°31’38.050” W) as a proxy for water level. Indeed, total daily rainfall was positively correlated to mean daily water level (collected during the period the logger was functional) at zero to five days later, but the Pearson correlation coefficient (*r*) computed with a 2-day lag on total daily rainfall exhibited the strongest association (*r* = 0.76; *P* < 0.01) compared to all other lag durations (0.25 < *r* < 0.55; 0.009 < *P* < 0.30) and was adopted as a predictor of detection probability in the analysis.

### Freshwater residence time analysis

The first step in our analyses was determining patterns in freshwater residence – the time elapsed between a fish’s release and first detection at the PIT array. To analyze freshwater residence time, we used a generalized linear model (GLM) with a generalized gamma distribution:

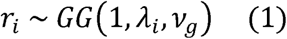

where *r_i_* is the freshwater residence time of fish *i*; *λ_i_* is the scale parameter for fish *i*; and *v_g_* is one of the two shape parameters of the distribution, which we allowed to vary by release group *g* (combination of release site and date for a total of nine groups). The other shape parameter was set to 1, so that the distribution reduces to the Weibull distribution (Plummer 2017). This approach implements the accelerated failure time parameterization of the Weibull distribution, which substantially improves mixing of chains during Bayesian implementation using JAGS compared to the use of the Weibull distribution proper in JAGS (Plummer 2017; see Model fitting and assessment). Using a log-link function, we modeled the scale parameter for an individual *i* as

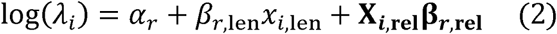

where *α_r_* is the intercept; *β_r_*_,len_ is the coefficient for the effect of an individual’s fork length (*x_i_*_,len_) at release; **β*_r_*_,rel_** is a vector of coefficients describing the main effects and two-way interactions between release site and date; and **X*_i_*_,rel_** is a design matrix with factor variables indicating the release site and date for an individual.

### Survival and detection analysis

To estimate outmigration survival, we used a state-space version of the Cormack-Jolly-Seber (CJS) capture-recapture model (Gimenez et al. 2007; Royle 2008), where survival and detection probabilities were constrained to be a function of covariates. The state-space formulation of the model is defined by two sub-models: an ecological state (alive or dead) sub-model describing the survival process and an observation (detected or undetected) sub-model describing the detection process of tagged individuals given their ecological state.

In the state space model, the ecological state process is defined as

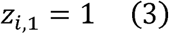

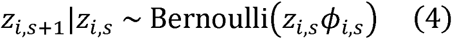

where *z_i_*_,1_ is the state of individual *i* at its release site, which is set to 1 as the individual was alive when released; *z_i,s_* and *z_i,s_*_+1_ are the states (0 = dead; 1 = alive) of individual *i* at site *s* and site *s* + 1 (release site: *s* = 1; upper antenna site: *s* = 2; and lower antenna site: *s* = 3); and *ϕ_i,s_* is the survival probability between sites *s* and *s* + 1 for individual *i.* We modeled individual survival probability from release site to the upstream PIT array (i.e. *ϕ_i_*_,1_) as

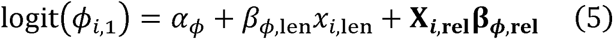

where *α_ϕ_* is the intercept; *β_ϕ_*_,len_ is the coefficient for the effect of an individual’s fork length (*x_i_*_,len_) at release; **β*_ϕ_*_,rel_** is a vector of coefficients describing the main effects and two-way interactions between release site and date; and **X*_i_*_,rel_** is a design matrix with factor variables indicating the release site and date for an individual.

During the initial model development, we assumed that survival probability for all fish sizes and release groups was 1 from the upper to the lower antenna (i.e. *ϕ_i_*_,2_) because of the short distance (100 m) between them. However, assessments of the model fit using posterior predictive checks (see Model fitting and assessment) were not satisfactory, but improved substantially after we removed the constraint and let the model estimate survival between antennas.

The observation process is conditional on the state of individual at a site and is defined as

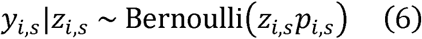

where *y_i,s_* is a variable indicating whether an individual *i* was detected (0 = undetected; 1 = detected) at a PIT antenna site (s = {2, 3}) given that it was alive and migrating past the site. We modeled individual detection probability at a PIT antenna site as

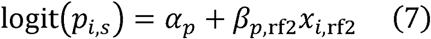

where *α_p_* is the intercept; *β_p_*_,rf2_ is the coefficient for the effect of rainfall two days before an individual migrated past the PIT array (*x_i_*_,rf2_) on detection probability, which is only known for individuals that were detected while migrating past the array. To assign a rainfall value for individuals that were not detected, we used the predicted date they would have migrated past the array had they survived. This date was calculated as the sum of their release date and the predicted number of days they would have resided in freshwater according to the model described in equation 2. Between May 25 and June 8, an error in the downstream antenna’s reader board prevented the detection of fish. For this reason, the detection probability of the downstream antenna was fixed at *p_i_*_,3_ =0 for fish that migrated past the array during that period (but detected at the upstream antenna) or for undetected fish whose migration date past the array was predicted to have occurred within the same period.

### Model fitting and assessment

The GLM of freshwater residence time and the state-space CJS model were fitted in an integrated framework using Bayesian inference (Kéry and Schaub 2012), which allowed full propagation of uncertainty in the parameters across the models (e.g. uncertainty in predictions of the day an undetected fish migrated past the PIT array and, therefore, the rainfall value that would influence its detection probability are automatically propagated within this integrated model fitting framework). We used Normal(0, 3.2) priors for the intercept and coefficients of all models described by eqs. 2, 5 and 7; Gamma(0.1, 0.1) hyperpriors for the shape and rate parameters of the prior Gamma distribution for the release-group-specific shape parameters *v_g_* in eq. 2; and a Beta(1, 1) prior for the survival probability between antennas (i.e. *ϕ_i_*_,2_).

The posterior distributions of model parameters were approximated with summaries of samples obtained with Markov Chain Monte Carlo (MCMC) algorithms implemented by JAGS 4.3.1 (Plummer 2017) via the package jagsUI 1.5.2 (Kellner 2021) in R 4.1.2 (R Core Team 2021). We ran four MCMC chains for 2,000,000 iterations, discarding the first 1,000,000 as burn-in and keeping every 1,000^th^ sample out of the remaining 1,000,000 samples, thus obtaining 1,000 samples per chain (total of 4000 samples). MCMC chains exhibited satisfactory convergence based on assessments using trace plots and their *R̂* statistic being < 1.1 (Gelman and Rubin 1992). The samples were used to summarize the posterior distributions of model parameters and predictions with their medians and 95% highest posterior density intervals (HPDI; McElreath 2020). For each coefficient in equations 2, 5, and 7, we also present the proportion of their posterior samples that had the same sign as their median (*f-*value, Plummer 2003). We considered *f* > 0.95 as indicative of strong evidence of an effect. Data summaries are presented as median, median absolute deviation (MAD), and ranges. Model predictions by release site and date were generated assuming an averaged sized fish in the dataset to control for the effects of size when comparing release strategies. Predictions are summarized with their median and 95% HPDI.

We assessed the fit of the freshwater residence model with posterior predictive check plots (Gabry et al. 2019), which indicated adequate fit (Figure S1 available in the Supplementary Material in the online version of this article). To assess the fit of the state-space CJS model, we compared the observed and expected frequencies of detection histories using a *χ*^2^ test. Detection histories were denoted by three digits (e.g. 101) referring to release (first digit) and detection at the two antennas (second and third digits for upstream and downstream antenna, respectively), where 1 indicates release/detection and 0 indicates no detection. To generate expected frequencies, we first simulated detection histories for each individual using parameter values sampled from their posterior distributions. Second, we computed the frequency of each detection history in a simulation. Lastly, we computed the median frequency of each detection history across 4000 simulations and used the median frequency of detection histories as the expected frequency in the *χ*^2^ test. The observed (100 = 3824; 110 = 432; 101 = 347; 111 = 242) and median expected (100 = 3811; 110 = 439; 101 = 364; 111 = 229) detection histories were similar, and the test results did not indicate lack of fit (*χ*^2^ = 12; d. f. = 9; P = 0.21; see also Figure S2).

## RESULTS

A total of 1,263 detections occurred during the monitoring period of May 23 to Aug 29, 1,021 of which were unique fish (21% of 4,845 released fish retained for analyses). Of these unique fish, 347 (34%) were detected exclusively on the downstream antennae, 432 (42.3%) were detected exclusively on the upstream antenna, and 242 (23.7%) were detected on both antennae. Furthermore, of the unique fish detected, 228 (22.3%), 398 (39%) and 395 (38.7%), were released in the lower river, lake, and upper river, respectively; whereas 179 (17.5%), 274 (26.8%), and 568 (55.6%) were released, respectively, on May 23, June 9, and June 19 (Table 1). Model estimates revealed that the PIT array had a baseline detection probability of about 0.4 under no rainfall (low water levels), which decreased rapidly with increased rainfall (and therefore high river levels), nearing zero above 20 mm rainfall events (Figure 2).

**FIGURE 2.**
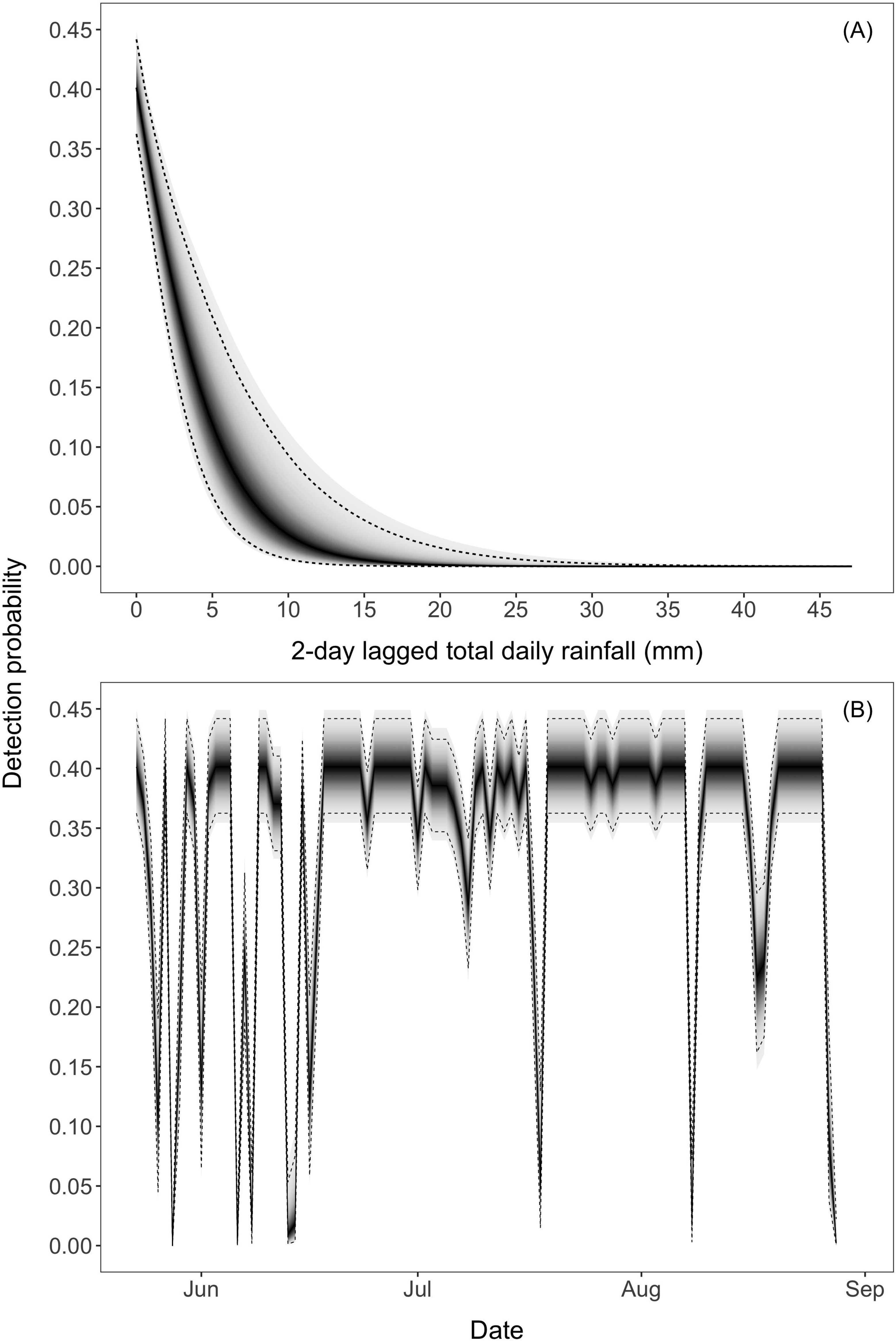
(A) Detection probability declined sharply with total rainfall being virtually zero at about total daily rainfall of 20 mm; and (B) high rainfall events were frequent during the first month of monitoring, resulting in several events of low detection probability. The solid line represents the median of the posterior distribution of detection probability at a given rainfall, whereas the dashed lines represent the 95% HPDI. The shading denotes the density of the full posterior distribution of detection probability, with higher probability densities denoted by darker shades. The median (95% HPDI) of the posterior distribution for the intercept was −1.03 (−1.33, - 0.72) and that for the effect of 2-day lagged rainfall was −1.90 (−2.69, −1.04).

The median freshwater residence time across all release sites and dates was 13.8 days and varied substantially within and between release groups (MAD: 9.1 days; range: 0.5 to 61.4 days; Table 2). Assessment of the posterior distribution of the freshwater residence model coefficients showed strong evidence for a negative relationship between fish length and freshwater residence time (*f* = 1, Figures 3A and 4), as well as for an interactive effect of release site and date on the time fish resided in freshwater (all interactions with *f* = 1; Figure 3A). The shape parameters of each release group (combination of release date and location) revealed that the smallest freshwater residence variation occurred among fish released in the lower river on May 23, followed by all the fish released on June 19 (Figure 3B).

**FIGURE 3.**
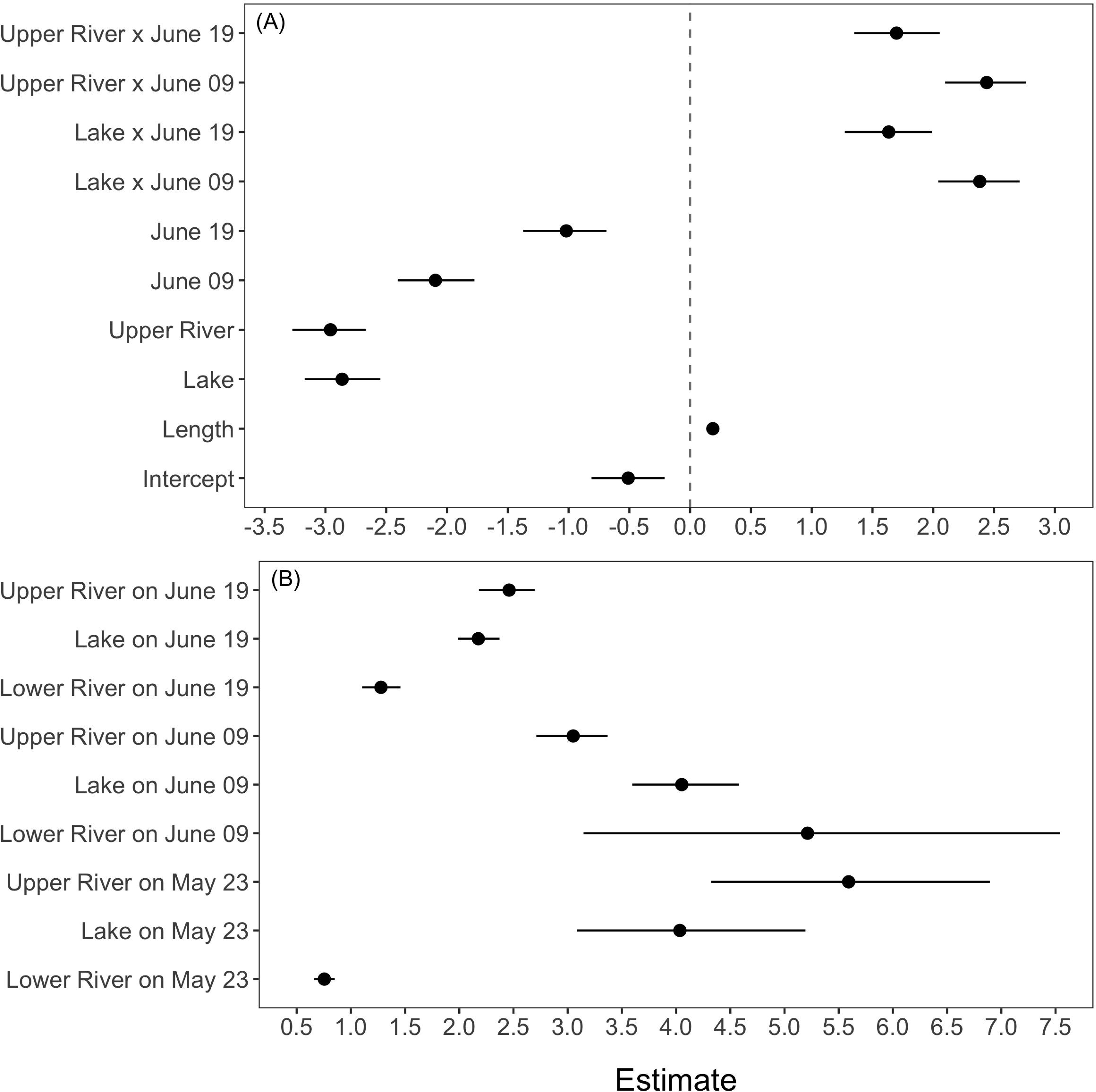
Medians (circle) and 95% HPDI (bars) of freshwater residence time model parameters. (A) Estimates of the intercept and coefficients for the effects of size at release, release site and date, and their interactions. The vertical dashed line indicates the value of zero. Note that the coefficients represent the effect on the scale parameter of the Weibull distribution, which is inversely related to its expected value (i.e. freshwater residence time). A positive effect on the scale parameter, therefore, translates into a negative effect on freshwater residence (decrease in time spent in freshwater). Lower river and May 23 are the reference levels for the release site and date factors and are represented by the model intercept. (B) Estimates of the shape parameters for release groups (combination of release site and date), where higher values indicate more variability in freshwater residence within a release group.

**FIGURE 4.**
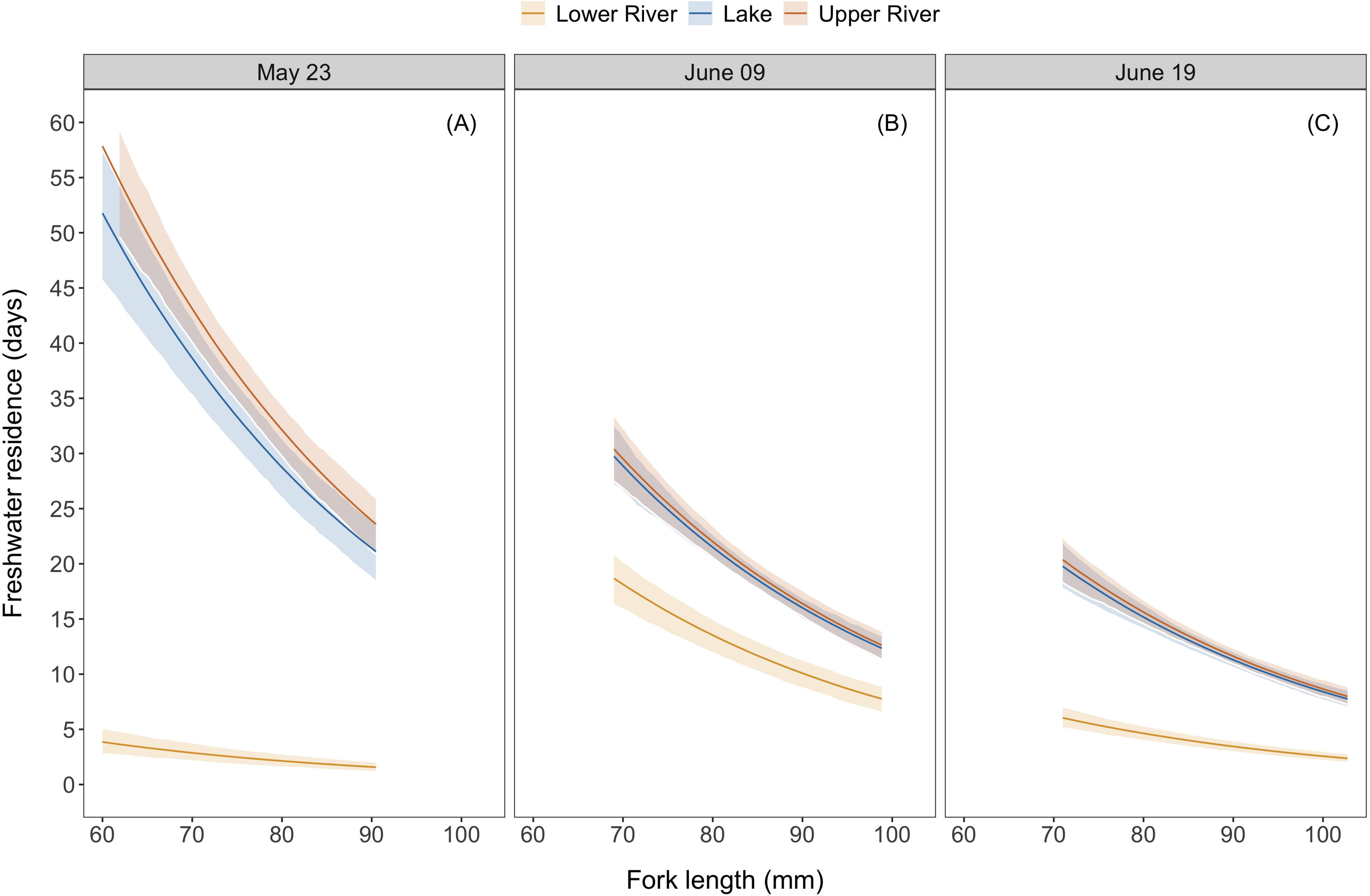
Predictions of freshwater residence time as a function of fish fork length (size) by release site and date. As fish length increases the freshwater residence time rapidly decreases. Predictions are shown for the range of sizes released on each date. Solid lines represent the median of the posterior distribution of the predictions and the ribbons represent the 95% HPDI.

Model predictions of residence times showed that fish released in the upper river on May 23 had the longest residence times at nearly 30 days followed closely by lake-released fish that same day (Figure 5A). Median freshwater residence time of both upper river and lake released fish decreased by ∼7 days with each subsequent release, ending at about 15 days for fish released on June 19 (Figures 5B and 5C). Fish released in the lower river exhibited a much lower and non-linear pattern of freshwater residence across release dates, being only a few days for fish released on the first (May 23) and last (June 19) release dates, and about 12 days for fish released in the intermediate (June 9) release date (Figure 5)

**FIGURE 5.**
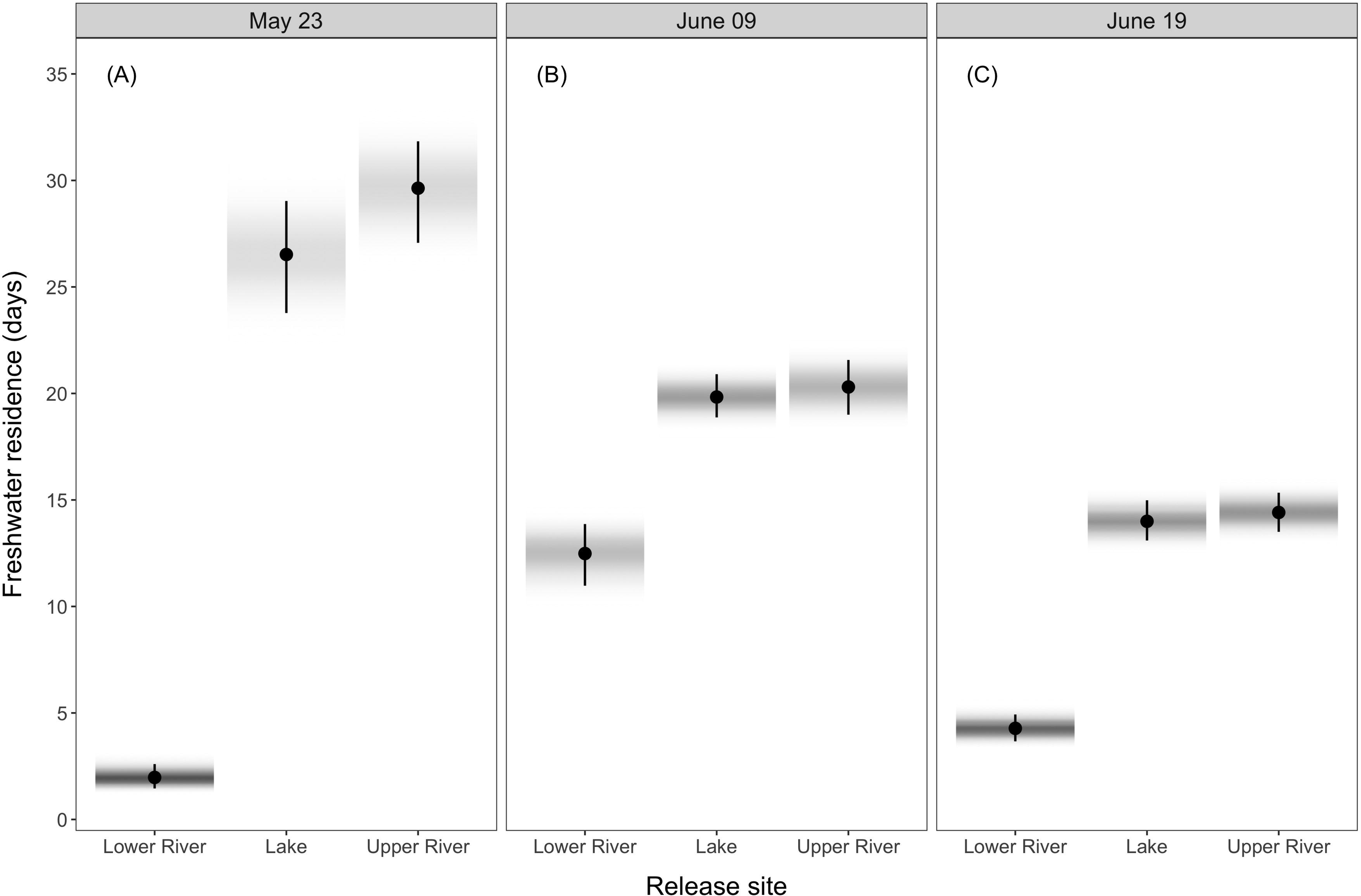
Predictions of freshwater residence time by release site and date. The further the fish were released from the estuary, the longer was their residence time. Regarding release date, fish released in the lake and upper river showed a decrease in residence time with each successive release date. In contrast, fish released in the lower river exhibited shorter residence times on the early release date, followed by an increase on the middle date before decreasing again with the final release. Predictions were made assuming a fish with length equal to the average in the data set (i.e. 82.7 mm). Circles represent the median of the posterior distribution of predictions and the bar shows their 95% HPDI. The shading represents the density of the posterior distributions of predictions (darker shades represent highest densities).

The median posterior prediction of survival probability across all release groups and fish sizes was 0.35 (95% HPDI: 0.02, 0.81). Similarly to the freshwater residence model, summaries of the posterior distribution of survival model coefficients revealed strong evidence for an interactive effect of release site and date on the probability of surviving the outmigration (all interactions had *f* = 1; Figure 6). Survival of upper-river- and lake-released fish were similar on all release dates, and both increased with later release dates (Figure 7). Fish released in the lower river had similar survival between the earliest (May 23) and latest release dates (June 19; Figures 7A and 7C), but exhibited low survival when released on June 9 (Figure 7B). Survival of lower-river-released fish was only higher than that of upper river and lake released fish on the earliest release date (Figure 7A). Survival probability between antennas was estimated as 0.89 (95% HPDI: 0.81, 0.96) for all release groups.

**FIGURE 6.**
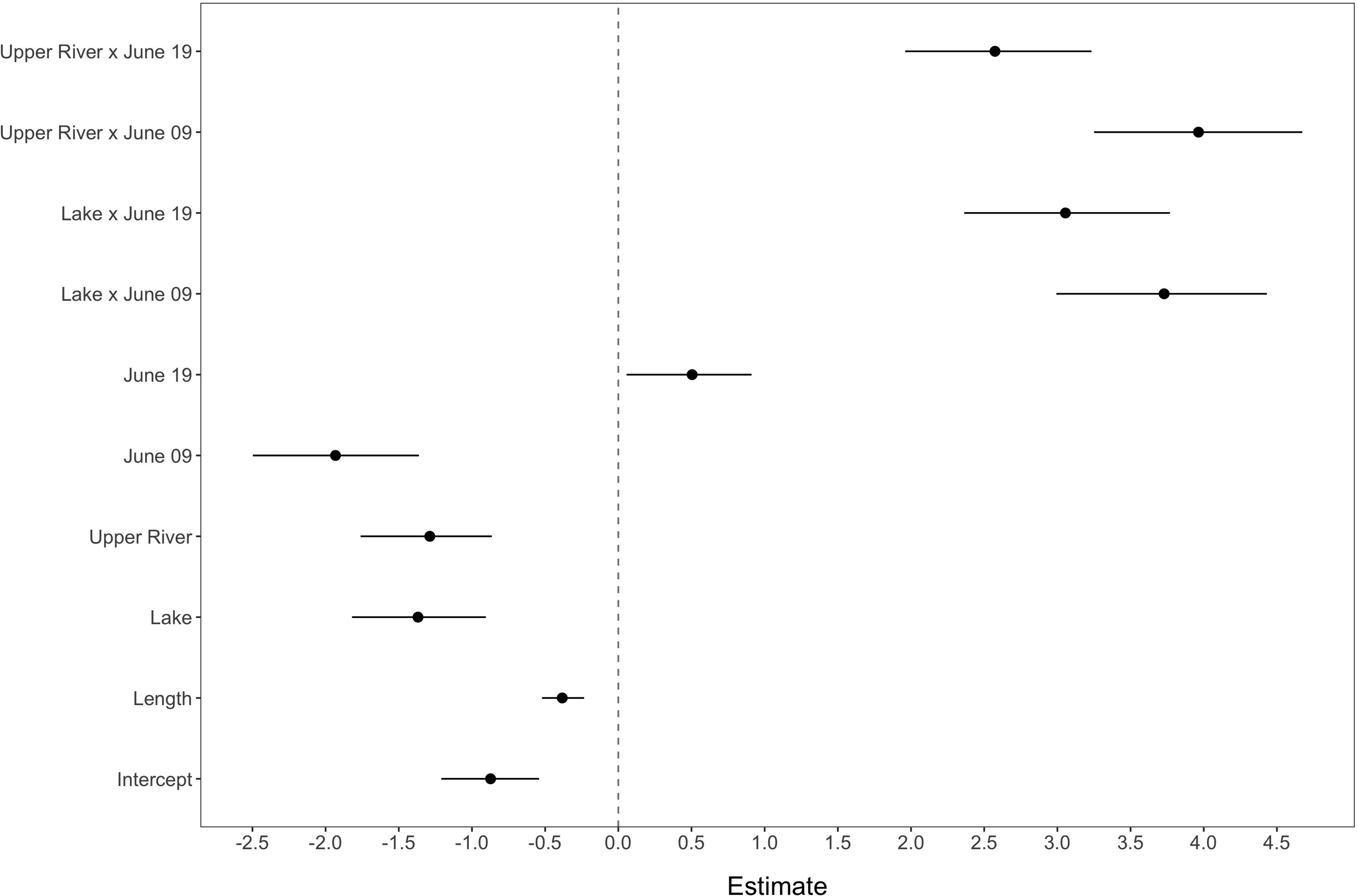
Medians (circle) and 95% HPDI (bars) of the survival probability model intercept and coefficients for the effects of size at release, release site and date, and their interactions. The vertical dashed line indicates the value of zero. Lower river and May 23 are the reference levels for the release site and date factors and are represented by the model intercept.

**FIGURE 7.**
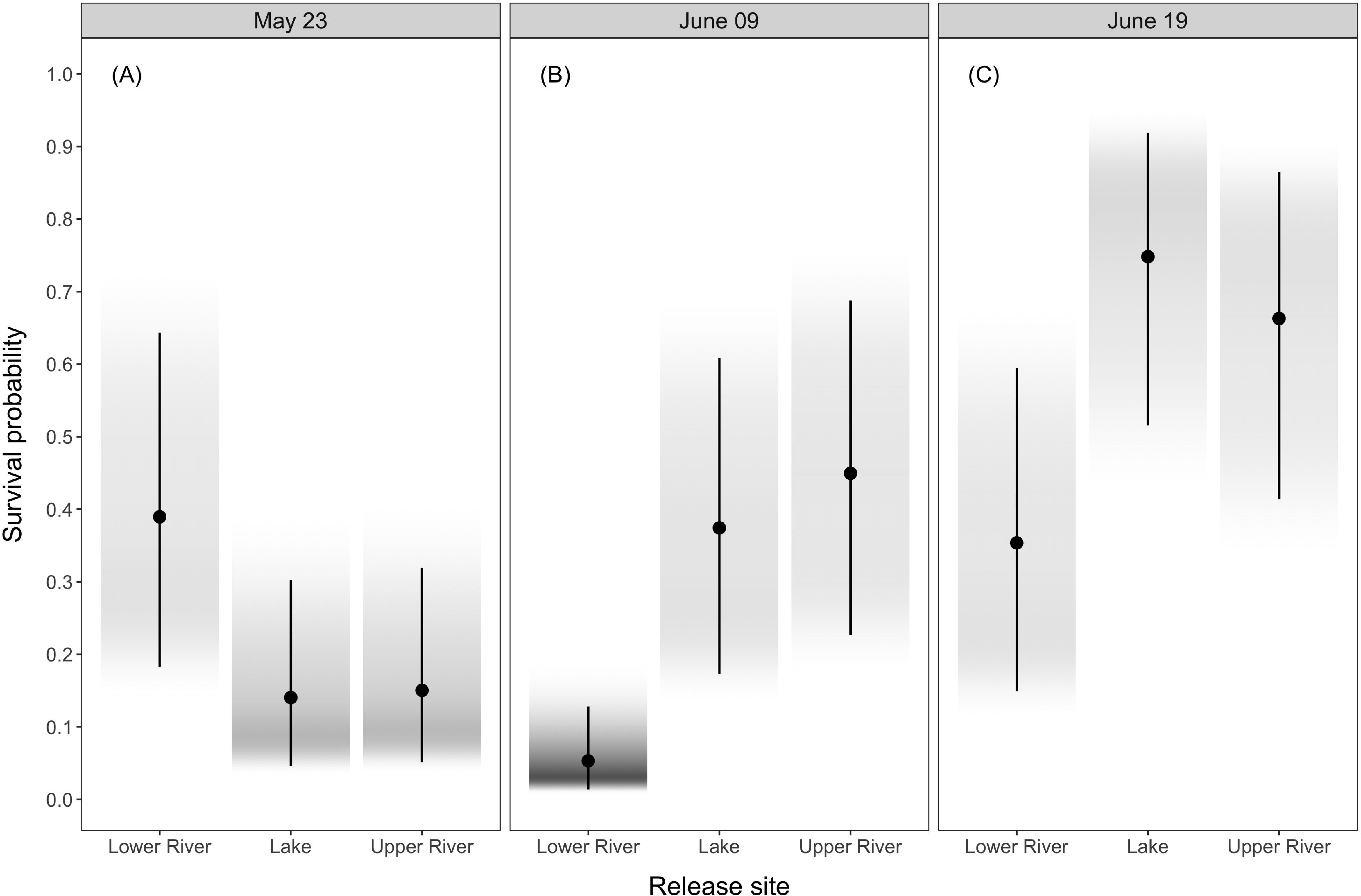
Predictions of survival probability by release site and date. Survival probabilities generally increased with later release for upper-river- and lake-released fish. For lower-river-released fish, survival probabilities were similar between the earlier and later releases and lowest for the intermediate (June 9) release. In general, uncertainty in predicted survival probabilities increased with later releases. Predictions were made assuming fish with length equal to the average in the data set (i.e. 84mm). Circles represent the median of the posterior distribution of predictions and the bar shows their 95% HPDI. The shading represents the density of the posterior distributions of predictions (darker shades represent highest densities).

When examining the relationship between length at release and detections, we observed a slight shift towards smaller fish being detected more frequently than larger fish (Figure 8). Indeed, the length at release of an individual fish had a negative effect on survival probability (Figure 9) and the *f-*value of 1 indicated strong evidence for this effect. Model predictions showed a marked decline in survival probability with size at release, with larger declines in survival occurring the higher the overall survival in a release group (Figure 9).

**FIGURE 8.**
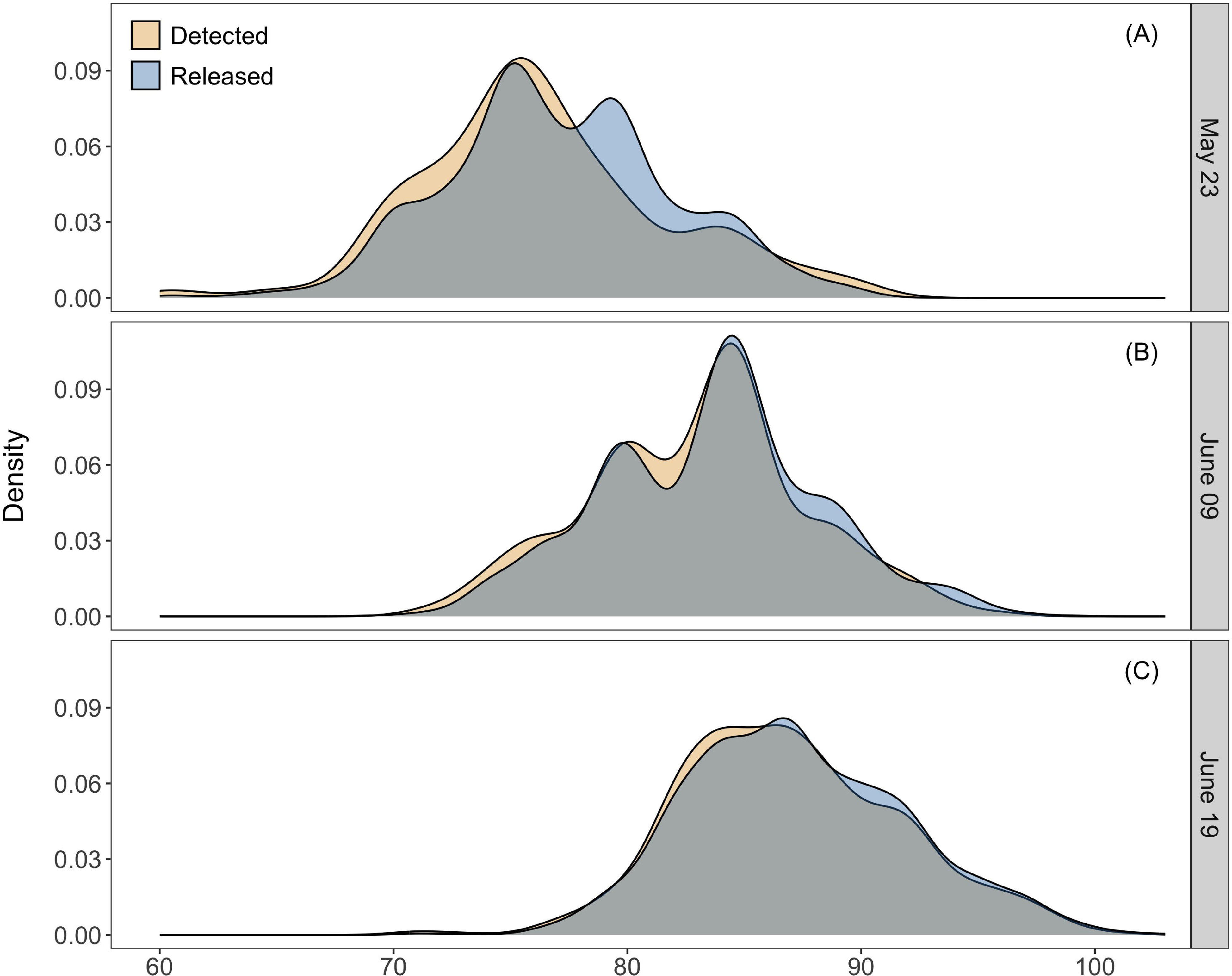
Probability density plots comparing the size distribution of fish at each release date to that of detected fish from the same release date. A consistent pattern emerges across all release groups revealing a slight skew towards smaller fish in the detected fish distribution.

**FIGURE 9.**
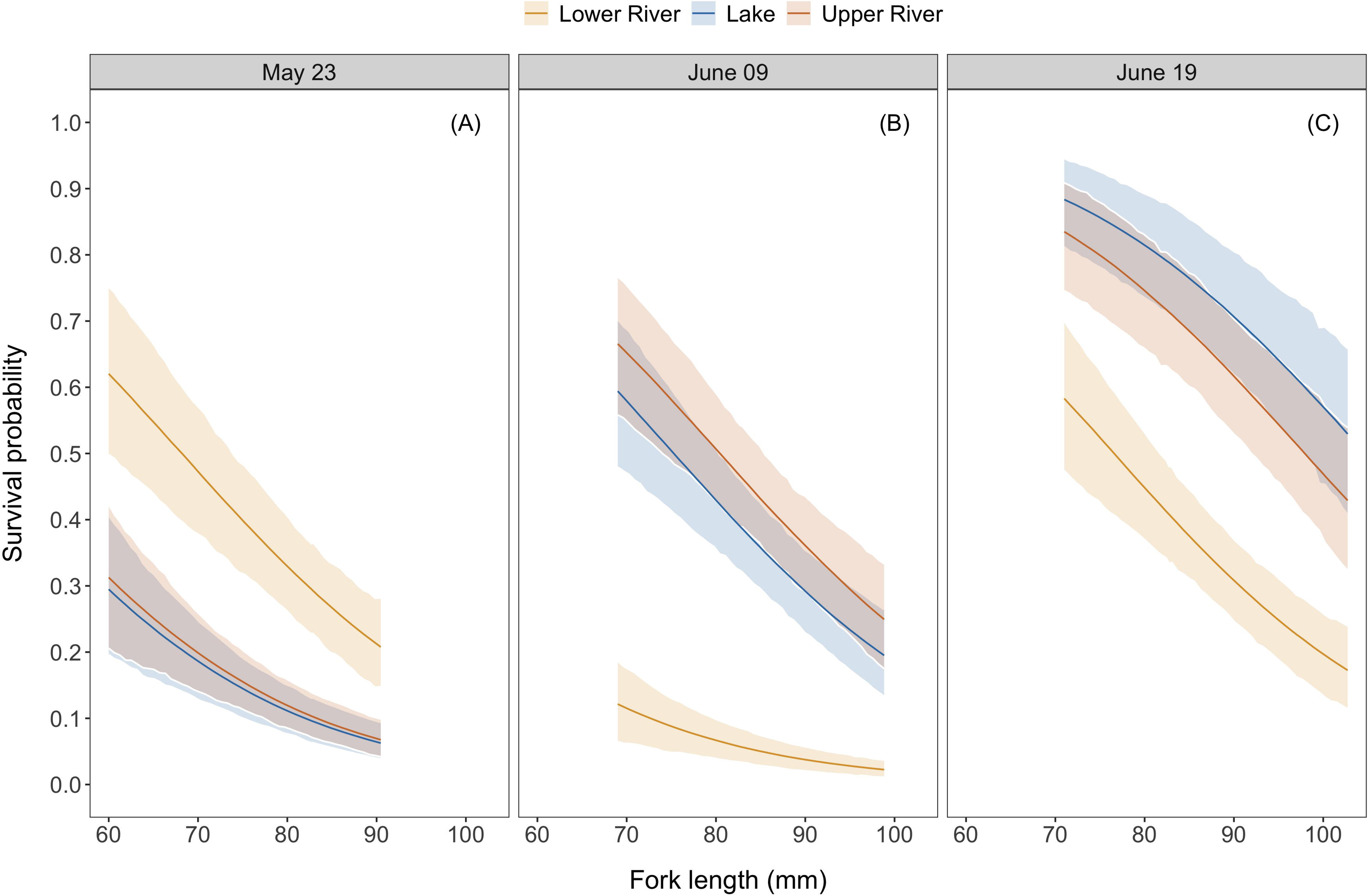
Predictions of survival probability as a function of fish fork length (size) by release site and date. Survival probability shows a marked decline as fish size increases. Predictions are shown for the range of sizes released on each date. Solid lines represent the median of the posterior distribution of the predictions and the ribbons represent the 95% HPDI.

## DISCUSSION

We used an integrated modeling approach and Bayesian inference to quantify freshwater residence and survival between freshwater release and estuary entrance of hatchery-origin Chinook Salmon smolts released in the Toquaht River at different sites and dates. We found that the time fish spent in freshwater post-release was highly variable, and that the site and date at which a fish was released strongly influenced residence duration. Survival probability was also highly variable across release sites and dates. The length of fish at release also influenced both freshwater residence time and survival, with smaller fish residing longer in freshwater and experiencing a higher probability of survival than larger fish. This work shows that hatchery release practices can markedly influence freshwater residence and survival and provide insight that managers can use to improve success of their programs.

The longer than expected freshwater residence time exhibited by several release groups and the marked negative relationship between size at release and residence time were surprising. Because of the short migration distance in the Toquaht River, and given that tagged fish were held in the hatchery until late June (well beyond the wild fish outmigration period of early March to late May) and grown to a size (4-10 g, and 70-100 mm) many times that of a wild fish of the same age (0.5–1 g and 40–50 mm), we expected that tagged fish would be ready to move quickly out of the river in a matter of hours to days from the release sites regardless of their size. Although some fish did outmigrate immediately after release, particularly those released in the lower river, other fish resided in the river for nearly two months, with smaller fish residing longer than larger fish. The range of residence time we observed could simply be hatchery fish expressing, like wild fish do, variation in freshwater residence strategies that are genetically predetermined or based on environmental conditions (Groot and Margolis 1991; Apgar 2021).

The early growth and outmigration cues in ocean-type Chinook Salmon, like the study population, are not dependant on photoperiod as in stream-type Chinook Salmon, and they do not experience growth or fitness reductions associated with staying in freshwater later into their outmigration period as seen in stream-type fish (Clarke et al. 1981). Without the photoperiod constraint on freshwater residence duration, the drivers of outmigration in ocean-type Chinook Salmon may be food, habitat, or water-quality related, wherein limitations on growth opportunities available in freshwater, and proximity to highly productive marine habitats govern freshwater residence duration and timing of marine entry (Levings et al. 1986; Chalifour et al. 2021). For the fish in this study, those that resided in freshwater longer may have done so to capitalize on feeding opportunities that could enable them to enter the marine environment with greater chances of surviving (Johnson et al. 2015). However, by residing in the river much longer than their wild counterparts, there is increased potential to be mismatched with food and habitat availability when they enter the estuarine and nearshore marine environment (Levings et al. 1986; Satterthwaite et al. 2014).

Despite the Toquaht River being a short (18 km), undammed river, the median survival probability for an average-size fish across all release groups was surprisingly low (0.35). This survival probability was somewhat higher than that for hatchery-origin Chinook Salmon outmigrating the San Joaquin and Sacramento Rivers (0.026-0.17; Michel 2019) in California and the nearby Cowichan River (0.24-0.64; Atkinson et al. 2024) on Vancouver Island. The Cowichan is the closest analogue to our study system but is over 40 km long and predator-dense compared to the Toquaht River. However, there are potential survival bottlenecks within the

Toquaht River, such as the lake outlet and base of the canyon, where predators are more concentrated and may have an outsized impact on juvenile survival (D. J. A. Hurwitz, personal observation), which is not without precedent. Chilko Lake Sockeye Salmon smolts, for example, exhibit very low survival within a few kilometers (0.32 over 13.5 km) at the start of their outmigration, which includes passage through the outlet of Chilko Lake, where Bull Trout *Salvelinus confluentus* congregate at high densities and prey on outmigrants (Furey et al. 2015; Rechisky et al. 2019). High mortality rates (37%) driven by predators over a short distance (3 km) immediately following release from a hatchery have also been observed in Coho Salmon released directly into the marine environment (Kanigan et al. 2024).

Accordingly, we expected that outmigrating Chinook Salmon released in the upper river and lake would experience high mortality at the outlet of Toquaht Lake, which is known to host the highest densities of piscivorous fish (e.g. Coastal Cutthroat Trout and Steelhead) in the river (D. Johnsen, Toquaht Lands and Resources Director, personal communication). Furthermore, we expected all release groups to experience high mortality at the base of the lower Toquaht River canyon, which also has high densities of predators (e.g. Common Merganser *Mergus merganser* and Short-Billed Gull *Larus brachyrhynchus*, adult Steelhead and Coastal Cutthroat Trout). However, despite lower-river-released fish being subject to potentially less predation pressure than fish released in the upper river and lake, we found that lower-river-released fish had substantially lower survival than fish released in the upper river and lake in two out of three release dates, suggesting that predation was not the sole and main factor affecting survival of release groups.

Periods of high and low survival aligned with the lowest and highest flow events, respectively, throughout the study. For example, between the first release on May 23 and May 27, rainfall and water levels remained low, and it was during this period that lower-river-released fish quickly outmigrated within the first 1–2 days and showed improved survival over upper-river- and lake-released fish. Heavy rain beginning on May 27 caused the water levels to rise significantly and remain high throughout the period in which lake and upper river fish released on May 23 would have been navigating the lower river canyon. The lower river canyon is the most tumultuous stretch of the river and fish migrating here during high flows are likely subject to injuries or disorientation (potentially increasing susceptibility to predation) (Elder et al. 2016). Indeed, the high rainfall and water level events between May 27 to June 16 coincided with the lowest estimates of survival probabilities for all release groups that had most fish outmigrating during that period (i.e. upper river and lake releases on May 23, lower river releases on June 9). In contrast, increases in survival probability were observed across all release groups that had most fish outmigrating after the high rainfall events (upper river and lake releases on June 9 and June 19, lower river releases on June 19). These findings are not an artefact of reduced detection probability during high rainfall events, as they were explicitly accounted for in our modeling of the observation process (but see below for another possibility when we discuss study limitations).

The potential relationship between low survival and high flows in our study differs from other studies showing that increased flows improve outmigration success and have a strong influence on overall cohort returns (Melnychuk et al. 2010; Buchanan et al. 2013; Michel et al. 2015; Cordoleani 2018; Michel 2019). However, these studies all occurred on dam-regulated California rivers, plagued by drought and where flows are heavily augmented. In these systems, increases in flows are linked to river conditions typically beneficial to salmon, such as increased velocity to speed migration, turbidity to hide juveniles from predators, and an overall increase in accessible habitat (Sommer et al. 2005). In contrast, the Toquaht River is not dam-controlled and has minimal land cover alteration or development. The river rises and falls rapidly, but rarely below a level that may harm fish or impede downstream migration. As a rainfall-driven system located in one of the rainiest parts of North America, the Toquaht River often experiences extreme rain events and flooding during the early rearing and outmigration period for salmon. Our findings suggest that increases in flows can reach a point where the benefits to fish are lost, and high flows in the Toquaht River can become a driver of mortality rather than a boost for survival probabilities.

Our finding that larger fish exhibit lower survival than smaller fish was surprising and contrasts with other studies that have found no relationship between size and survival of outmigrating Chinook Salmon (Michel et al. 2015; Pellet et al. 2019). One potential explanation for our results is that larger fish were mismatched to forage in the habitats that the Toquaht River provides, leading to either faster seaward migration or lower survival for those remaining in freshwater. This explanation is based on work conducted in the estuary showing that smaller wild fish were able to utilize a wider range of habitats and surpass the initially larger hatchery fish in size (Levings et al. 1986). Alternatively this finding may be a result of size selective predation, wherein predators such as harbour seals and birds preferentially target the larger fish (Nelson et al. 2019; Thomas et al. 2017). Another potential explanation is the propensity for larger fish to take up a precocious parr or mature river parr life history (Bernier et al. 1993; Johnson et al. 2012), in which they never leave freshwater and spawn the following fall. That is, the decline in survival with size that we observed may not have been mortality, but rather larger fish choosing to reside in freshwater indefinitely, or at least beyond the four-month study period. These fish would not be at risk of being detected (as they would not migrate past the PIT array) and are equivalent to tagged animals that permanently emigrate from study areas in classical capture-recapture studies, leading to negative biases in survival (Nichols 2006).

One of the fundamental limitations of this study, and of many tag-based studies of juvenile fish, is working within the restrictions of minimum tag sizes. The minimum fork length for 12 mm tags was set at 69 mm (after initially using 60mm), which is already large in comparison to wild West Coast of Vancouver Island Chinook Salmon, and required holding fish in the hatchery until later in the season to allow them time to grow to the minimum tagging size. This limitation prevented the creation of a release group closer in size and timing to wild fish. The reason for using 12 mm tag size speaks to another prominent challenge in working with PIT tags: the tradeoff between tag size and read-range. Modern PIT tag detection systems using 12 mm FDX tags can commonly achieve read ranges of 35–41cm (Connolly et al. 2008; Beeman et al 2012). This makes them highly effective in shallow water, confined spaces, and flow-controlled rivers, but limits their utility in medium-sized and larger rivers, and rivers with highly variable water levels. The antennas used in this study were able to achieve a vertical read range of 20–25 cm over a 14 m span of the river channel, resulting in a detection probability ranging from 0-0.40. Reducing the length of our cable would have achieved a higher vertical read range but reduced the width of river channel each antenna covered. Under baseflow conditions, our array effectively covered the channel horizontally and vertically. However, as rainfall and water levels increased, larger portions of the river channel, both horizontally and vertically, became uncovered by the detection array. Consequently, fish swimming by the uncovered areas would be equivalent to tagged animals that permanently emigrate from the study area in classical capture-recapture study designs, and cannot be distinguished from mortalities, leading to negative biases in survival (Nichols 2006). This may explain in part the low survival estimated for fish outmigrating during high flows. The survival estimate of 0.89 in the short distance (100m) between antennas may also be associated with incomplete coverage of the river during high flows, suggesting that on average about 11% of the fish migrating through the lower antenna would not be at risk of detection if our expectation of 100% survival between antennas were true.

Another limitation of this study is the high potential for tag collision, which can occur when a large group of tagged fish passes over the antennas, overwhelming the equipment’s capacity to accurately detect each individual fish (Zentner et al. 2021). As outmigrating salmon tend to move in schools (Hoar 1958), and since our study fish were released all at once, they likely passed over the antennas in large groups, potentially leading to tag collisions, lower detection probabilities than we would estimate based on rainfall (water levels) alone and, consequently, reduced precision of our survival estimates (Nichols 2006).

Assessments of hatchery release strategies present valuable opportunities for managers to refine their programs and meet their production and/or conservation objectives. However, there is a notable lack of detailed understanding regarding how these strategies specifically influence freshwater residence and outmigration survival. Our research shows that both freshwater residence and outmigration survival can vary substantially with size at release and strategies based on the site and date of releases. In our study system, the findings suggest that releasing small fish later in the season and in the lake and upper river would maximize the number of fish that survive to reach the Toquaht River estuary. Even though we observed increased freshwater survival with smaller fish and later releases, it has been shown that entering the estuary earlier and at a larger size can result in better survival in the marine environment and results in more returning adults (James et al. 2023). However, maximizing returns may not be the only hatchery objective and fish released at a larger size also tend to return as smaller and younger adults than those released at a smaller size (Feldhaus et al. 2016). Therefore, future assessments of release strategies should evaluate whether increases in outmigration survival afforded by a given strategy carry over to the early marine life or are offset by a potential mismatch with marine conditions. Additionally, it is important to assess how factors beyond survival (e.g. age and size at return) are influenced by release strategies. Finally, streamflow may be more important than the site or date of release in influencing outmigration survival in streams similar to the Toquaht River, and efforts should be made to not expose newly released fish to high flow events. Traditionally, Thornton Creek Hatchery’s releases have been based on the assumption that predation would be lower at high flow levels and that timing releases with increased flows would improve survival. Our results suggest that this practice needs to be re-evaluated as survival of outmigrating fish is likely to be higher if the fish do not encounter high flows in the lower river canyon. Although our results may be applicable to other similar systems, the unique and complex dynamics involved in each enhancement program require research that is focused on specific facilities and objectives, while a changing climate requires continual re-evaluation of hatchery practices and an informed and adaptive approach to future salmon management (James et al. 2023). Recognizing the immense diversity of rivers and salmon populations, and that there is no one size fits all approach, carrying out projects in understudied regions is an important step in the future of salmon enhancement and recovery planning.

## ACKNOWLEDGEMENTS

This project was funded by the Toquaht Nation Government, Mitacs, Clayoquot Biosphere Trust, Pacific Salmon Foundation, and Natural Sciences and Engineering Research Council of Canada (through a Discovery Grant to EGM). In-kind support was provided by the Department of Fisheries and Oceans Canada, the British Columbia Conservation Foundation, Thornton Creek Salmon Enhancement Society, and Redd Fish Restoration Society. We thank the Toquaht Nation for welcoming us into their territory and trusting us to carry out this work. We would also like to thank our field team Emily Fulton, James Costello, Miles Downsbrough, Collin Middleton, Christian Carson and Kaylyn Kwasnecha who were instrumental in executing this project. We want to thank Drs. Mark Shrimpton, Erin Rechisky, and Cameron Freshwater for their suggestions during the study and the review of this work. Dr. Scott Hinch provided helpful comments on our manuscript. Our research was enabled in part by support provided by the University of Waterloo’s Graham computing system and the Digital Research Alliance of Canada (<u>alliancecan.ca</u>).

## ETHICS STATEMENT

This research meets the ethical guidelines and legal requirements of Canada and followed the guidelines of the Canadian Council on Animal Care. Fish handling and tagging procedures were approved by the University of Northern British Columbia Animal Care and Use Committee (protocol no. 2021-04) and carried out under a Fisheries and Oceans Canada Aquaculture licence AQSEP 129556.

## DATA AVAILABILITY STATEMENT

Data collected in this study as well as code used to process and analyze the data and model output are available at https://datadryad.org/stash/share/aMvY_YidTVFE5LydNHaunyr8-f-L924ZHQmNDQaKswU (private for peer review) and later at https://doi.org/10.5061/dryad.6t1g1jx6t

