## Supplementary Material for "Release strategies affect the freshwater residence and survival of hatchery-reared juvenile Chinook Salmon"

Supplementary Material: Posterior Predictive Checks

### Freshwater residence model

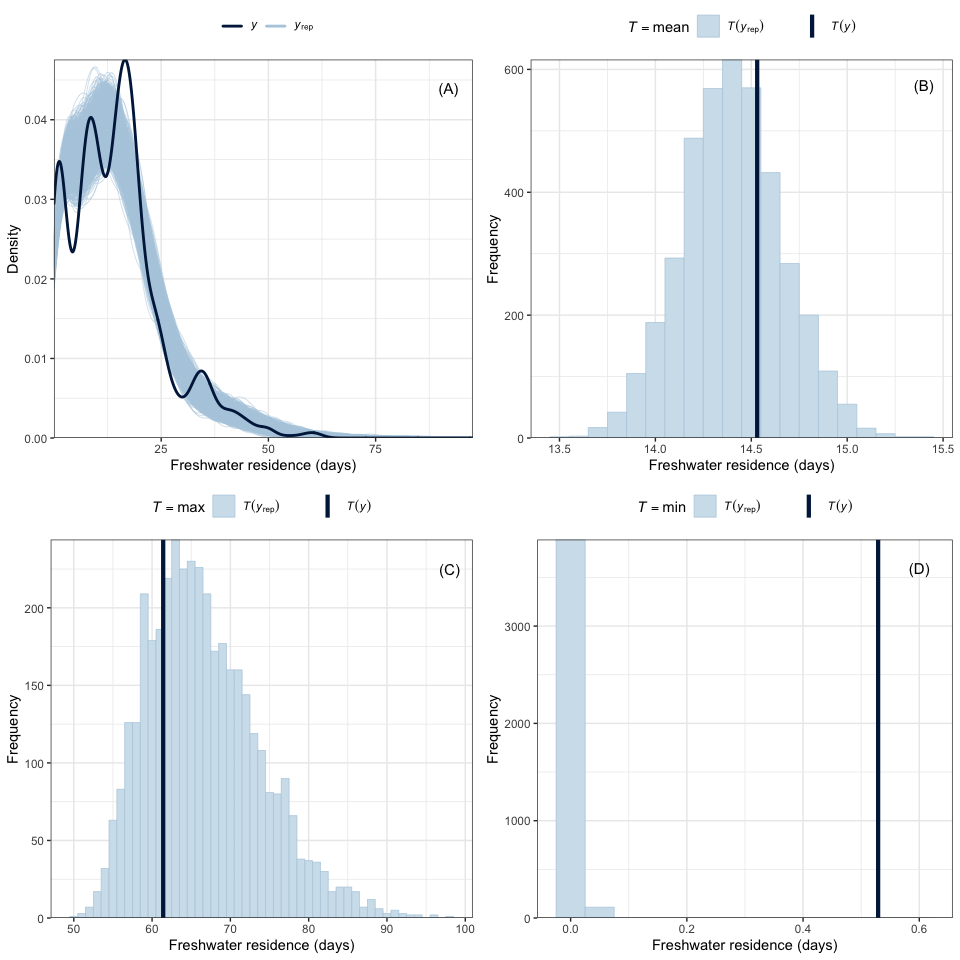

**FIGURE S1** Posterior predictive plots show that the (A) distribution of model predictions of freshwater residence ($y_{\text{rep}}$, light blue) generally included the observed distribution of freshwater residence times ($y$, dark blue) in the dataset. The distribution of the (B) mean and (C) maximum values of posterior predictions also included the observed mean and maximum freshwater residence times. However, the distribution of (D) minimum posterior predictions of freshwater residence did not include the observed minimum freshwater residence time in the dataset, though the difference was only < 1 day.

### State-space CJS model

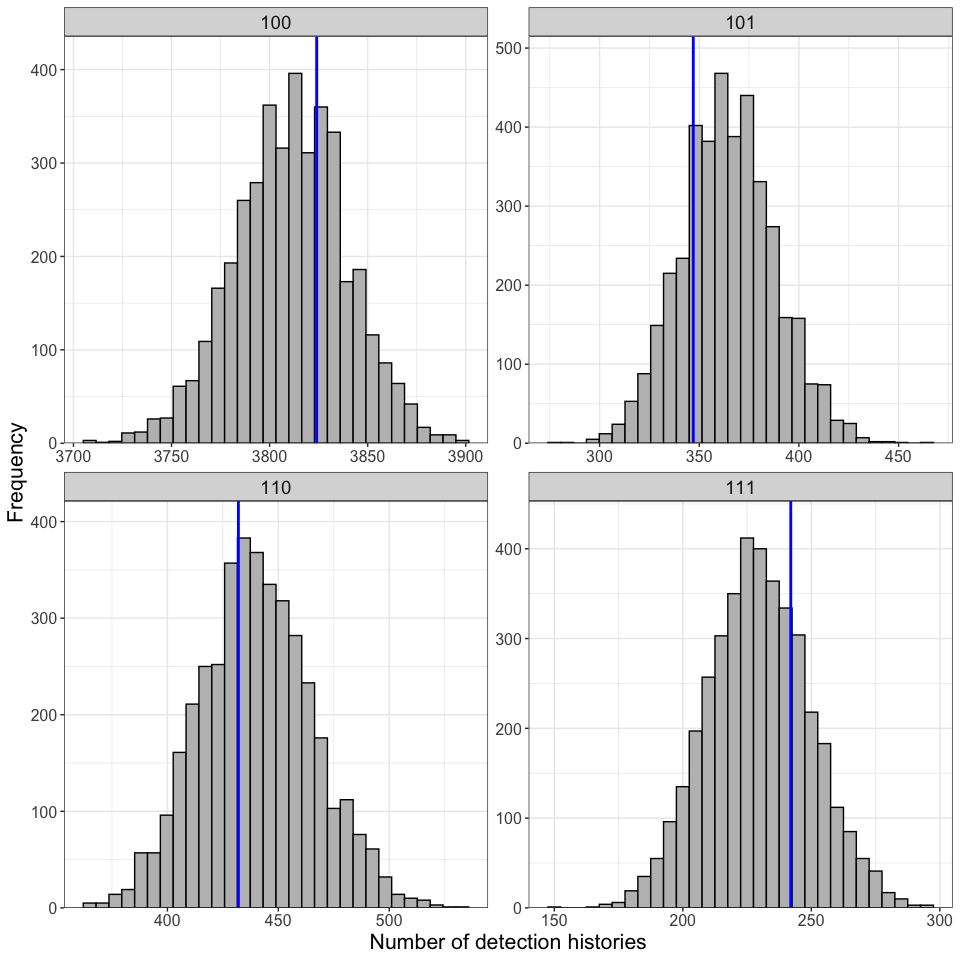

**FIGURE S2** The distribution of posterior replicates of detection histories included the observed number of each detection history in the study (vertical blue lines). Detection histories show at the strip on top of plots are represented by three digits with the first one denoting release (1). The two subsequent digits denote detection (1) or non-detection (0) at the upper and lower antenna sites.
